# A new rat model of creatine transporter deficiency reveals behavioral disorder and altered brain metabolism

**DOI:** 10.1101/2020.08.11.246330

**Authors:** Lara Duran-Trio, Gabriella Fernandes-Pires, Dunja Simicic, Jocelyn Grosse, Clothilde Roux, Pierre-Alain Binz, Carmen Sandi, Cristina Cudalbu, Olivier Braissant

## Abstract

Creatine is an organic compound used as fast phosphate energy buffer to recycle ATP, important in tissues with high energy demand such as muscle or brain. Creatine is taken from the diet or endogenously synthetized by the enzymes AGAT and GAMT, and specifically taken up by the transporter SLC6A8. Deficit in the endogenous synthesis or in the transport leads to Cerebral Creatine Deficiency Syndromes (CCDS). CCDS are characterized by brain creatine deficiency, intellectual disability with severe speech delay, behavioral troubles such as attention deficits and/or autistic features, and epilepsy. Among CCDS, the X-linked creatine transporter deficiency (CTD) is the most prevalent with no efficient treatment so far. Different mouse models of CTD were generated by doing long deletions in the *Slc6a8* gene showing reduced brain creatine and cognitive deficiencies or impaired motor function. We present a new knock-in (KI) rat model of CTD holding an identical point mutation found in patients with reported lack of transporter activity. KI males showed brain creatine deficiency, increased urinary creatine/creatinine ratio, cognitive deficiency and autistic features. *Slc6a8*^*xY389C*^ KI rat fairly enriches the spectrum of CTD models and provides new data about the pathology, being the first animal model of CTD carrying a point mutation.

## INTRODUCTION

Creatine (Cr) is a nitrogenous organic acid used as phosphate energy buffer to recycle ATP by the Cr-PCr system; Cr is also one of the main osmolytes of the brain, and suggested to play a role as neuromodulator or neurotransmitter ^1,2^. Cr is taken from the diet or endogenously synthetized in a two-step pathway by the enzymes AGAT (arginine-glycine amidinotransferase) and GAMT (guanidinoacetate methyltransferase), and specifically transported into the cells by SLC6A8 (also called CRT, CreaT or CT1, henceforth CrT) ^2^. Guanidinoacetate (GAA) is synthetized from arginine (Arg) and glycine (Gly) in the first step, and it is methylated generating Cr in the second step. Cr breaks down spontaneously in creatinine (Crn).

Deficits in endogenous synthesis or transport of Cr cause Cerebral Creatine Deficiency Syndromes (CCDS) ^3-6^. CCDS are characterized by brain Cr deficiency, intellectual disability with severe speech delay, behavioral abnormalities such as attention deficits and/or autistic features, and seizures for GAMT and SLC6A8 deficiencies ^7^. Among CCDS, Cr transporter deficiency (CTD) is caused by loss-of-function mutations in the *SLC6A8* gene (for a scheme of the human locus with the position of the mutations, see **Figure 1A**). CTD is a X-linked gene disorder with an estimated prevalence of about 2% of males with intellectual disability with, unlike the other two CCDS, no efficient treatment so far ^1,7,8^.

**Figure 1:**
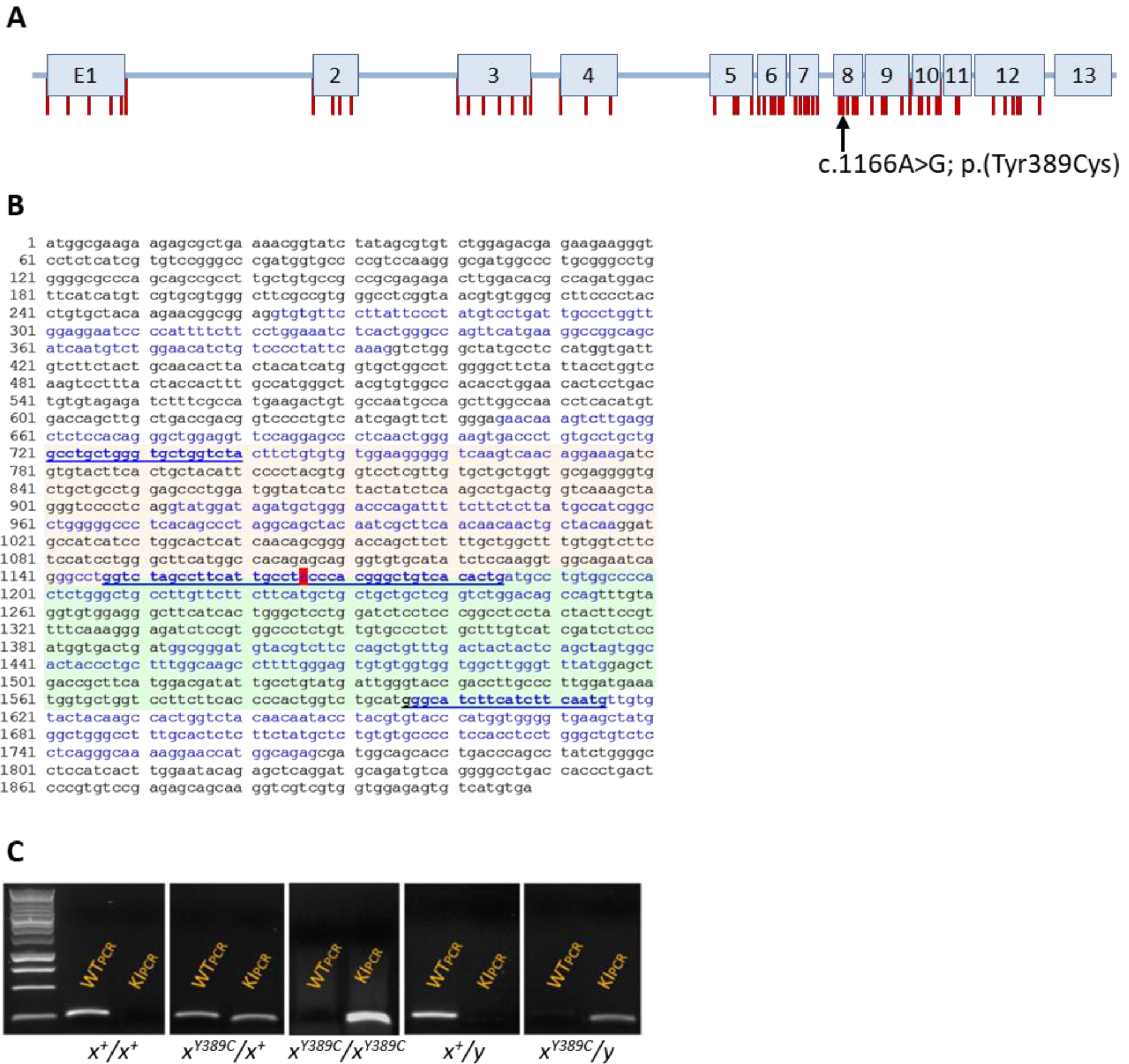
Generation and genotyping of Slc6a8Y389C rat. (**A**) Scheme of human SLC6A8 locus with the exons and the position of mutations found in patients (vertical red bars; missense mutations or little deletions according to Van de Kamp et al. 2013). Black arrow points to the position of the mutation c.1166A>G. (**B**) Wild-type rat cDNA sequence showing the alternating exons (black and blue letters) and the position of the mutation (red highlight). Green and orange backgrounds represent WT and KI fragments, respectively, after PCR amplification. Underlined letters are the sequences related to the primers used. (**C**) PCR products for each genotype. WT fragment at the left, KI at the right.

To better understand the pathology and with the ultimate goal of finding a treatment, several mouse models for CTD have been generated ^9-16^. In particular, CrT knock-out (KO) mouse males with ubiquitous deletions of the exons 2-4 ^9,16^, 5-7 ^13,14^ and 5-8 ^12^ showed brain creatine deficiency and decreased body weight, features present in CTD patients ^17^. Ubiquitous CrT KO males with exons 5-8 deleted showed impaired motor function ^12^. Cognitive deficits with normal sociability were demonstrated in the ubiquitous KO males with deletions in exons 2-4 ^9,16^ and 5-7 ^13,14^. Deletion in exons 2-4 also caused some changes in monoamine neurotransmitters correlating with higher resilience to stress, while deletion in exons 5-7 leaded to more stereotypies, impaired working memory performance and some brain alterations associated with aging.

Brain-specific KO mice of CrT have shown additional and somehow different phenotypes. Deletion of exons 2-4 ^11^ or 5-7 ^13,18^ in CNS nestin-positive cells exhibited brain Cr deficiency but normal Cr levels in periphery. Brain CrT KO of exons 2-4 mice maintained the same learning and memory deficits as the corresponding ubiquitous KO, plus hyperactivity and normal body weight ^11^. In contrast, brain CrT KO of exons 5-7 mice showed a rescue in the general phenotype (no stereotypies, normal memory performance and body weight) but declined at early age, correlating with some brain alterations related with aging ^13,18^. Deletion of CrT exons 2-4 in CaMKII-positive cells (excitatory neurons) ^15^ revealed similar phenotype as the nestin-targeted KO mouse, while targeting only DAT-positive cells (dopaminergic neurons) exhibited hyperactivity ^19^.

All the accumulating knowledge using these various mouse models highlights the importance of the Cr transporter in brain function and physiology, and allows a better understanding of SLC6A8 deficiency. However, CTD patients do not always present the same phenotype ^17^. Further studies are thus needed to fully understand CTD pathology, and to allow, through the use of various complementary animal models, the best translation to human pathology ^20^.

As an alternative to mouse models, rats have a number of potential advantages over mice. In particular, they are closer to human physiology, biochemistry and behavior in comparison with mice ^21,22^. In order to increase the fundamental knowledge and to increase the relevance of preclinical studies with complementary preclinical models, we generated a novel rat model of CTD: the *Slc6a8*^*Y389C*^ knock-in (KI) rat line. *Slc6a8*^*Y389C*^ KI rat holds a point mutation found in CTD patients which was reported to completely abolish Cr transporter activity ^7^. Here we show the first analysis of *Slc6a8*^*xY389C/y*^ rat KI males as model of CTD, being the first KI animal model of CTD.

## METHODS

### Generation, genotyping and housing of the *Slc6a8*^*Y389C*^ KI rat strain

Based on the patient *SLC6A8* c.1166A>G; p.(Tyr389Cys) point missense mutation reported as completely abolishing Cr transporter activity ^17^ (**Figure 1 A**), we designed a KI *Slc6a8* c.1166A>G; p.(Tyr389Cys) rat line (henceforth *Slc6a8*^*Y389C*^, **Figure 1 B**). As in human, the rat *Slc6a8* gene is located on the X chromosome, thus 3 *Slc6a8*^*xY389C/y*^ male (KI) and 3 heterozygous *Slc6a8*^*xY389C/x*^ female founders were generated from the Sprague-Dawley (OFA-SD) rat strain using CRISPR/Cas9 technology (Cyagen Biosciences Inc, USA). *Slc6a8*^*Y389C*^ rats were viable and fertile, and crossbreedings between wild-type males (WT) or KI males and wild type females or heterozygous females generated litters with the expected Mendelian ratios and with the 5 possible genotypes: (WT and *Slc6a8*^*xY389C/y*^ KI males; WT, heterozygous *Slc6a8*^*xY389C/x*^ and homozygous *Slc6a8*^*xY389C/xY389C*^ females; **Figure 1 C**). For the present study, crossbreedings between WT males and heterozygous females were made to obtain WT and *Slc6a8*^*xY389C/y*^ KI males in the same litter.

Genotyping was standardized using PCR amplification (QIAGEN kit ref. 201445). DNA was obtained from ear punch using the KAPA BIOSYSTEMS kit (ref. KK7352) and finally diluted in nuclease-free water. Amplification primers were: for WT fragment (250bp) F-WT/5’-GGTCTAGCCTTCATTGCCTA-3’ and R-WT/5’-GCCCTCCACACCTACAAACT-3’; for KI fragment (211 bp) F-KI/5’-ACTCTGGCATGAGACCCTGT-3’ and R-KI/5’-CAGTGTGACAGCCCGTGGGC-3’. PCR conditions were as follows: 10min 94°C, 2x[30s 94°C, 30s 67°C, 30s 72°C], 36x[30s 94°C, 30s 66°C, 30s 72°C], 10min 72°C, store at 4°C. A representative PCR of the 5 possible genotypes is shown in **Figure 1 C**.

Animals were maintained under a 12h/12h light-dark cycle. Food and water were available *ad libitum*. All experiments were performed with the approval of the veterinary authorities of the Canton de Vaud (Switzerland; authorization VD-3284) and in accordance with the regulations of the Swiss Academy of Medical Science. Efforts were made to minimize stress and number of animals used.

### Proton magnetic resonance spectroscopy (^1^H-MRS) in CNS at high magnetic field (9.4T)

For ^1^H-MRS 3 months-old rats were anesthetized with isoflurane (4% during induction; 2% for maintenance) using a nose mask under 70% compressed air and 30% O_2_. Body temperature (kept at 37.5 ± 1.0 °C using circulating warm water) and respiration were monitored continuously by a small-animal monitor system (SA Instruments, New York, USA). In vivo ^1^H-MRS was performed on a horizontal actively shielded 9.4 Tesla system (Magnex Scientific, Oxford, UK) interfaced to a Varian Direct Drive console (Palo Alto, USA). A home built ^1^H-quadrature surface coil (14 mm diameter) was used as transceiver and placed over the head of the animal. The volumes of interest (VOI) were positioned in dorsal hippocampus (2×2.8×2 mm^3^, 4 WT and 5 KI), striatum (2.5×2×2.5 mm^3^, 4 WT and 5 KI) and cerebellum (2.5×2.5×2.5 mm^3^, 3 WT and 4 KI) using axial and sagittal anatomical T_2_ weighted images (multislice turbo-spin-echo sequence, repetition time TR=4 s, effective echo time TE_eff_=52 ms, echo train length=8, field of view=23×23 mm^2^, slice thickness=1 mm, 2 averages, 256×256 image matrix). The static magnetic field homogeneity was adjusted using first and second order shims by FAST(EST)MAP leading to water linewidths of 10-12Hz for hippocampus and striatum and 14-16Hz for cerebellum ^23^. Localized spectra were acquired with SPECIAL spectroscopy sequence (TE=2.8 ms, TR=4 s, 160 averages) to obtain the specific neurochemical profile in the 3 mentioned VOIs ^24,25^. Metabolite concentrations were estimated using LCModel software ^26^ and water as internal reference. The LCModel basis-set for spectral fitting contained a spectrum of macromolecules acquired *in vivo* ^27^ and 17 individual metabolites measured *in vitro*: alanine (Ala), ascorbate (Asc), aspartate (Asp), glycerophosphocholine (GPC), phosphocholine (PCho), creatine (Cr), phosphocreatine (PCr), γ-aminobutyrate (GABA), glutamine (Gln), glutamate (Glu), glutathione (GSH), myo-inositol (Ins), lactate (Lac), N-acetylaspartate (NAA), N-acetylaspartylglutamate (NAAG), phosphoethanolamine (PE) and taurine (Tau). PCho and GPC were expressed as tCho (PCho+GPC) due to better accuracy in the estimation of their concentration as a sum. The ultra-short echo-time ^1^H-MRS allowed the detection of the abovementioned 17 metabolites ^24^.

### CSF, urine and plasma collection

The same day of the sacrifices, 6 WT and 7 KI rats aged of 3 months were anesthetized with 4% isoflurane under 70% compressed air and 30% O2. Fur around the base of the head and the dorsal part of the neck was shaved with an electric shaver. A little incision was made in medial position and longitudinally through the skin and muscles at the base of the skull. The cavity was maintained opened using little separators and cleaned of blood using cotton swabs. Carefully, puncture in cisterna magna was made to collect CSF. After CSF collection, the rat was positioned in supine position and the abdominal cavity was opened to collect urine from the bladder. Finally, the thoracic cavity was opened and blood was collected transcardially with heparin (5000 u/ml) and centrifuged at 2710 x g for 7 min. Supernatant (plasma) was transferred in a new tube. Liquids (CSF, urine and plasma) were stored at −80 °C for further analysis.

### Measure of creatine, guanidinoacetate and creatinine in plasma, urine and CSF

Cr and GAA were quantified by liquid chromatography coupled to mass spectrometry (LC-MS) following a μSPE sample preparation protocol. For plasma and urine, 180 μl of a 0.28 μM aqueous solution of internal standards (d_3_-Creatine, Sigma-Aldrich and ^13^C_2_-Guanidino acetic acid, VU Medical Center Amsterdam) containing 2.2% formic acid (v/v) was added to 20 μl of centrifuged plasma or urine. For CSF, 100 μl of 0.5 μM internal standard solution was mixed with 100 μl of CSF sample. The samples were loaded on a MCX μSPE plate (Oasis® MCX μElution Plate 30 μm, Waters Corp.) previously activated with MeOH and water, and then washed with formic acid 2% (v/v) and MeOH. Elution was obtained with 0.67 mol/l NH_4_OH solution on collection wells prefilled with acetonitrile. After centrifugation, samples were directly injected onto a LC-MS system (Waters I-Class UPLC system coupled with a Waters XEVO TQS micro mass spectrometer). Separation of the analytes was performed in a 4.5 min gradient at a flow rate of 0.5 ml/min on a HILIC column (ACQUITY UPLC^®^ BEH HILIC, 2.1 × 100 mm, 1.7 μm, Waters Corp.) maintained at 30 °C, with a 0.2% formic acid containing ammonium formiate (10 mmol/l):acetonitrile mobile phase gradient. Mass spectrometry was performed by electrospray ionization in positive mode using two SRM (Selected Reaction Monitoring) transitions (one quantifier and one qualifier) for each analyte. The m/z values of the selected quantitative and qualitative transitions, respectively, were the following: 118 → 100.85 and 118 → 30 for GAA, 120 → 103 and 120 → 31 for 13C2-GAA, 132.1 → 43.82 and 132.1 → 90 for Cr, 135 → 46.88 and 135 → 93 for d3-Cr. Quantified concentrations were calculated with TargetLynx (Waters Corp.). Creatinine (Crn) was measured in urine by a COBAS 8000 automate (Roche, Switzerland).

### Behavioral tests

Two cohorts of males were used to do the behavioral tests: grooming and Y-maze for spatial memory test in one; novel object recognition (NOR), Y-maze for spontaneous alternation and Social preference/Social memory (SP-SM) tests for the other one. WT and KI were littermates in each cohort and the set of tests began at 12 weeks of age.

#### Grooming

Rats were introduced in a round (1m diameter) open arena with black walls and floor made of black PVC for 10 min while spontaneous behavior was recorded with a videocamera. Grooming behavior was scored blindly with the Observer XT® software and cumulative duration and frequency of such behavior were analyzed. 10 WT and 10 KI males were used.

#### Y-maze for spatial memory test

A Y-shape maze of 50 cm arm length, grey walls and floor made of grey PVC in conditions of 4 lux in each arm was used. For the learning phase, the rat was introduced in the Y-maze for 10 min with one arm closed. After 4 hours, the rat was located in the same starting arm and was allowed to explore all the arms for 5 min. EthoVision® software was used to track the time spent in each arm. Exploration time (tN + tF, in seconds) and Recognition Index ((tN - tF)/(tN + tF)) were calculated from the cumulative time spent in the familiar (tF) and the new (tN) arms. 9 animals per group were used.

#### Novel Object Recognition (NOR) test

An arena of 57×36cm with grey walls and floor made of grey PVC in 4 lux conditions was used. For habituation, the rat explored the empty arena for 5 min. On next day, the rat was introduced in the arena with two identical objects for 5 min in the learning phase. After 4 hours, the rat was allowed to explore again the arena for 5 min but one of the objects was changed for a different one. Both, new and familiar objects and the right and the left sides were randomized. All phases were recorded with a videocamera and exploratory behavior was scored blindly with the Observer TX® software. The cumulative duration times for exploring familiar (tF) and new object (tN) in testing phase were used to calculate Exploration Time (in seconds, tN + tF) and Recognition Index ((tN – tF)/(tN + tF)). 8 WT and 9 KI males were used.

#### Y-maze spontaneous alternation

The rat was introduced in the abovementioned Y-maze and allowed to explore the three arms opened for 8 min. EthoVision® software was used to track rat position and calculate the number of total entries in all the arms (n° entries in arm A + arm B + arm C) as well as the percentage of alternation (100*frequency of alternations/(n° total entries - 2)). 8 WT and 10 KI males were used.

#### Social Preference and Social Memory (SP-SM) tests

A three-chamber arena 80×34cm with grey walls and floor made of grey PVC with 10 lux in the center was used for SP-SM test. A transparent cylinder made of PVC, 13 cm in diameter and with openings, was positioned in each lateral chamber. During habituation, a juvenile rat was located in one cylinder while an object was positioned in the other one, and the experimental male rat was allowed to explore only the center chamber for 5 min. During the social preference phase, doors were opened and the rat could explore all the chambers (with the juvenile rat or the object) for 10 min. After 2 hours, the object was changed for another juvenile, and the experimental rat was allowed to explore both new and familiar juveniles for 10 min. Relative positions of the items (object and juveniles) were randomized. All phases were recorded with a videocamera and exploration behavior towards the object or the juveniles was scored blindly with the Observer®. Cumulative duration time exploring each item, the number of visits (frequency) per each item, and the first item to be explored in each phase were taken into account for the analysis. 9 WT and 10 KI were used.

### Statistical analysis and graphs

All statistical analysis and graphs were conducted with R-3.5.1 ^28^. A Shapiro test was used to assess the normality of each sample. To address the significance between WT and KI groups or between each group with respect to the value expected by chance, Mann-Whitney tests were performed when normality was rejected while, for normal distributions, we used Bartlett tests to evaluate equality of variances and t-tests for equality of the means.

To calculate the log_10_ of the ratio KI/WT concentrations of different metabolites in brain, a pseudocount (+0.001) was added to all concentrations before doing the ratio. Graphs were done using ggplot2 package ^29^, while heatmaps were generated with gplots ^30^.

In the analysis of first decision for the SP-SM experiment, we evaluated different questions. First, we asked if there was a significant bias in the decision of exploring first one of the two possibilities. We modelled this question considering a Bernouilli variable with probability q=0.5 (no preference), and estimating the probability of selecting one of the two choices with a binomial distribution of parameters q, N, k, where N is the number of animals tested and k the number of visits of one of the items. We tested if the preferred choice was significantly different than random performing a two-tail exact binomial test, or a one-tail exact binomial test when prior information existed (e.g. when a given behavior was observed for the WT and the preference of the KI was tested, or when the preference was known a priori from literature). Finally, we tested if the preferences of KI and WT were significantly different between them using a Barnard’s CSM Test (exact.test, method=CSM, package Exact ^31^).

## RESULTS

### Generation of the *Slc6a8*^*Y389C*^ rat

To obtain a model as near as possible of the human CTD, a knock-in (KI) rat was generated with the *SLC6A8* c.1166A>G; p.(Tyr389Cys) missense mutation found in human patients and reported with no functional activity of the transporter ^17^ (**Figure 1**). The obtained *Slc6a8*^*Y389C*^ rats were viable and bred normally, generating litters with the expected Mendelian ratios.

### *Slc6a8*^*Y389C*^ rats are deficient in brain Cr

CTD patients are characterized by CNS Cr deficiency and are diagnosed, as all CCDS patients, using brain ^1^H-MRS. High magnetic field 9.4T ^1^H-MRS spectra in hippocampus (**Figure 2 A**) as well as other brain regions (striatum, cerebellum; data not shown) systematically showed much lower peaks corresponding to Cr and PCr in *Slc6a8*^*xY389C/y*^ KI males compared to those of WT ones. Concentration of Cr and PCr in *Slc6a8*^*xY389C/y*^ KI males was 70-75% less than that of WT in every brain region that was measured (**Figure 2 B**). These measures by ^1^H-MRS validate our *Slc6a8*^*Y389C*^ rat model as brain Cr deficient and therefore as a valuable model to study CTD.

**Figure 2:**
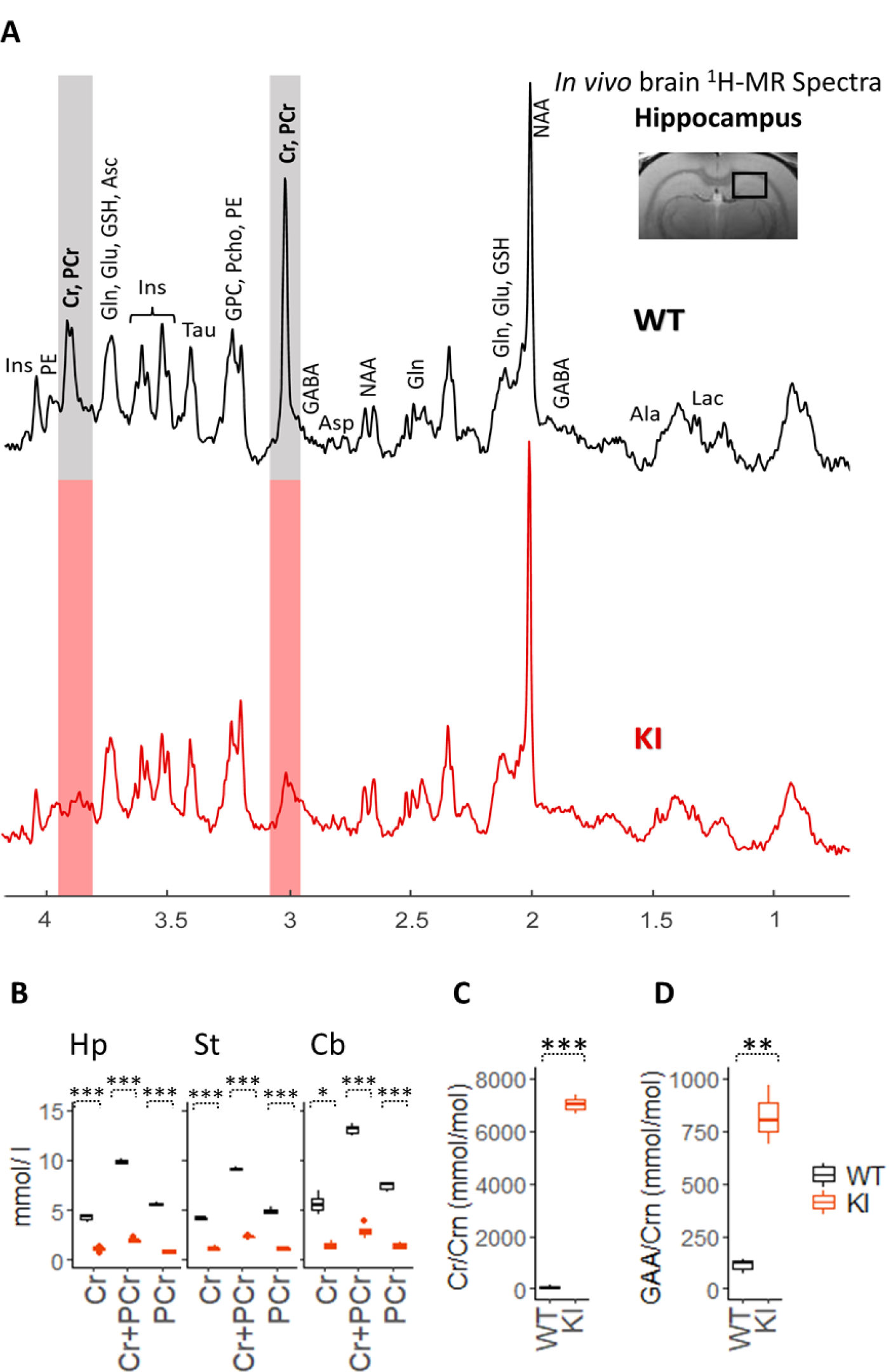
Brain creatine deficiency and renal creatine transporter deficiency in *Slc6a8*^*xY389C/y*^ KI males. (**A**) Representative 9.4T ^1^H-MRS spectra in the hippocampus of WT and *Slc6a8*^*Y389C*^ KI rats, showing the strong decrease of the Cr and PCr peaks in *Slc6a8*^*Y389C*^ KI rats. Note the much smaller Cr and PCr peaks in KI male. The localization of the measured voxel is presented on top of the panel. (**B**) Box plots of Cr and PCr concentrations in hippocampus (Hp), striatum (St) and cerebellum (Cb) in WT and *Slc6a8*^*Y389C*^ KI males (black and orange boxplots, respectively). 3-4 WT and 4-5 KI animals. (**C**) Cr/Crn and (**D**) GAA/Crn ratios in the urine of WT and *Slc6a8*^*xY389C/y*^ KI males. 3 animals per group. * : p<0.05, **: p<0.01, ***: p<0.001 (two-tail t-test).

### Lack of functional creatine transporter in *Slc6a8*^*Y389C*^ rat males

CTD patients show an increased Cr/Crn ratio in urine due to the deficiency in functional Cr transporter present on kidney proximal tubular cells (which normally allows the reuptake of Cr from primary urine). We thus measured Cr and Crn in the urine of our *Slc6a8*^*Y389C*^ rats to further validate our model as Cr transporter-deficient. *Slc6a8*^*xY389C/y*^ KI males showed a very strong significant urinary increase of Cr/Crn ratio (6999 versus 56 mmol/mol, p<0.001, **Figure 2 C)**, thus validating the loss of function of the Cr transporter in our *Slc6a8*^*Y389C*^ model. This increase is due to an increase of Cr (12163 versus 521 μmol/l, p=0.055) and a significant decrease of Crn (1.7 versus 8.5 mmol/l, p=0.005). We also measured GAA in urine, and show, very interestingly, that GAA/Crn ratio was also strongly and significantly elevated in *Slc6a8*^*xY389C/y*^ KI males (817 versus 112 mmol/mol, p<0.001; **Figure 2 D**), although no significant changes were found in urinary GAA levels (1502 versus 949 μmol/l, p=0.50).

### *Slc6a8*^*Y389C*^ rat males present a decrease of Cr and an increase of GAA levels in plasma

We collected plasma and CSF from WT and *Slc6a8*^*xY389C/y*^ KI males at around 3 months of age to see whether Cr and GAA concentrations were altered. We show a significant decrease of Cr and a significant increase of GAA concentrations in plasma of KI males in comparison to those of WT males (**Figure 3A** left and right panels, respectively). Cr and GAA levels in CSF from *Slc6a8*^*xY389C/y*^ KI males at around 3 months of age showed no differences in comparison to those of WT males (**Figure 3B** left and right panels, respectively).

**Figure 3:**
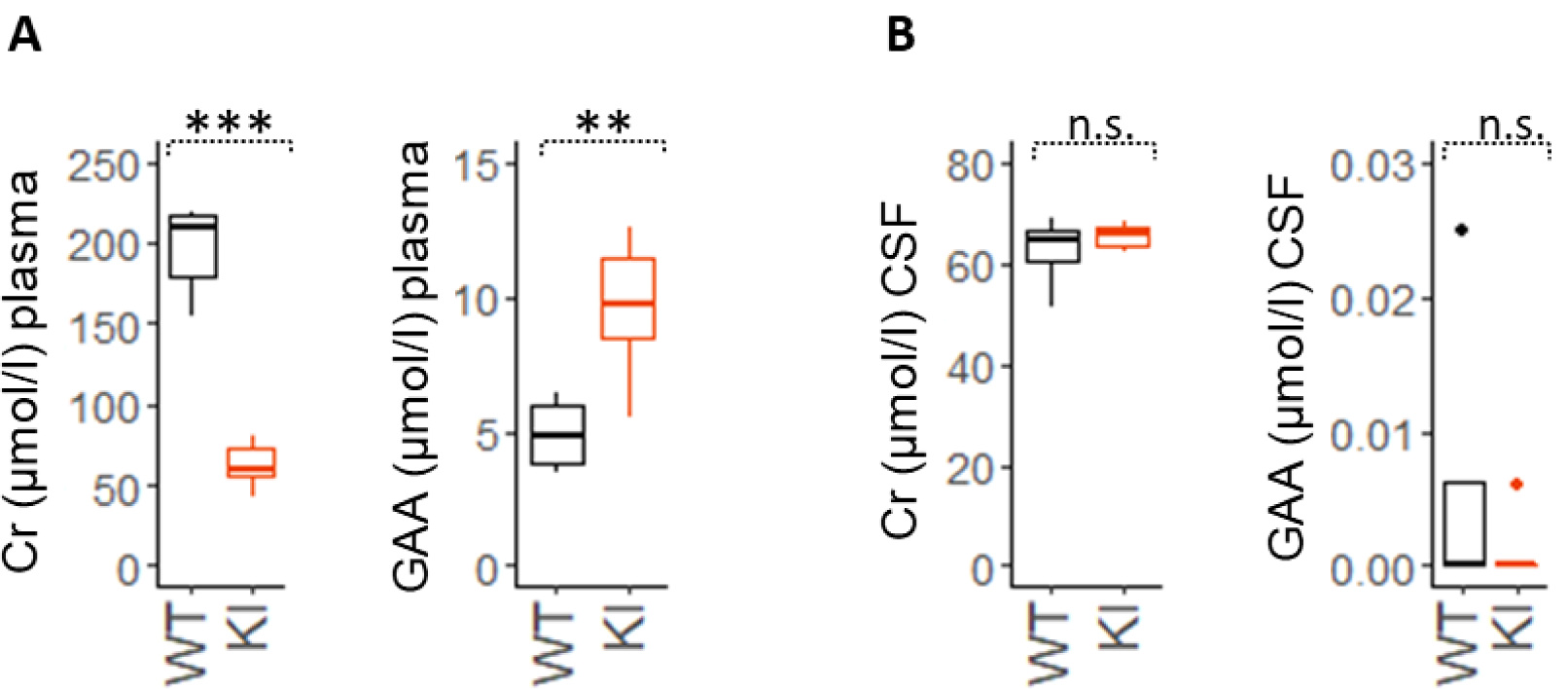
Cr deficiency and GAA increase in plasma, while no change in CSF, in *Slc6a8*^*Y389C*^ rats. (**A**) Box plot of Cr and GAA concentrations (left and right panel, respectively) of plasma in WT (black) and KI (orange) males. Note the reduction in Cr and the increase in GAA levels. (**B**) Box plot of Cr and GAA concentrations (left and right panel, respectively) of CSF in WT and KI males. 6 WT and 7 KI males; **: p<0.01, ***: p<0.001, n.s. : no statistical differences (two-tail t-test).

### *Slc6a8*^*Y389C*^ rat weight gain is slowed

KI males (*Slc6a8*^*xY389C/y*^) presented a significant reduced body weight gain just after weaning, as compared to WT males, showing for instance 56% of the body weight of their WT littermates at 14 weeks of age (312.7 ± 8.6 g *versus* 524.6 ± 12.1 g, p<0.001, Mann-Whitney test) (**Figure 4**).

**Figure 4:**
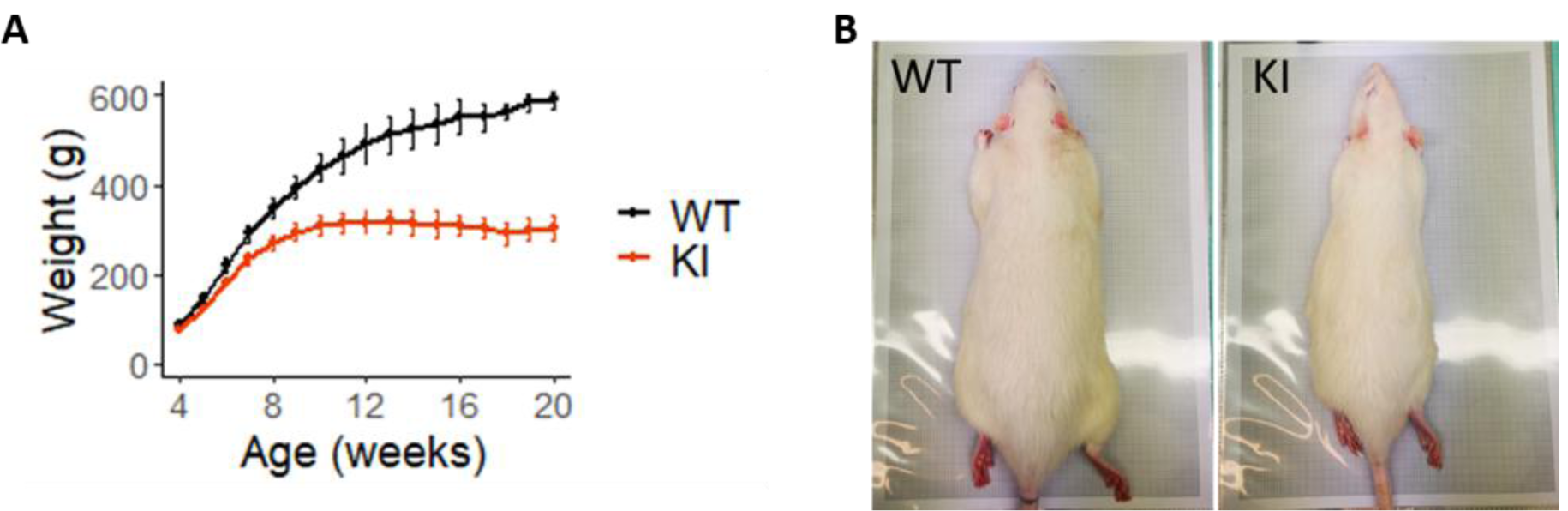
*Slc6a8*^*Y389C*^ KI males show a significantly decreased body weight gain. (**A**) Weight average (in grams) from 4 to 17 weeks of age. Error bars represent standard deviation. 13 animals per group, p<0.0025 from 4 weeks-old on (Mann-Whitney test). (**B**) Body size comparison between a WT and a KI male at 20 weeks of age. Gray rectangle dimension is 280×190 mm.

### Other brain metabolic changes in *Slc6a8*^*Y389C*^ rats

As described above, *Slc6a8*^*Y389C*^ rats showed brain Cr deficiency demonstrated by ^1^H-MRS in every CNS region measured (**Figure 2A-B**). We further took advantage of our high resolution 9.4T ^1^H-MRS measures to quantify potential metabolic changes in the brain of *Slc6a8*^*xY389C/y*^ KI males. Interestingly, Gln, NAA, and the sum of NAA and NAAG (NAA+NAAG) concentrations were significantly increased in all the brain regions analyzed while other metabolites changed their levels only in specific brain regions (**Figure 5 A and B**). In hippocampus, GABA, Glu, Tau and total choline (GPC+PCho) were significantly increased. In striatum, Ins increased while Lac decreased significantly. Interestingly, Ins tended to change in all brain regions although in different directions (**Figure 5 B**).

**Figure 5:**
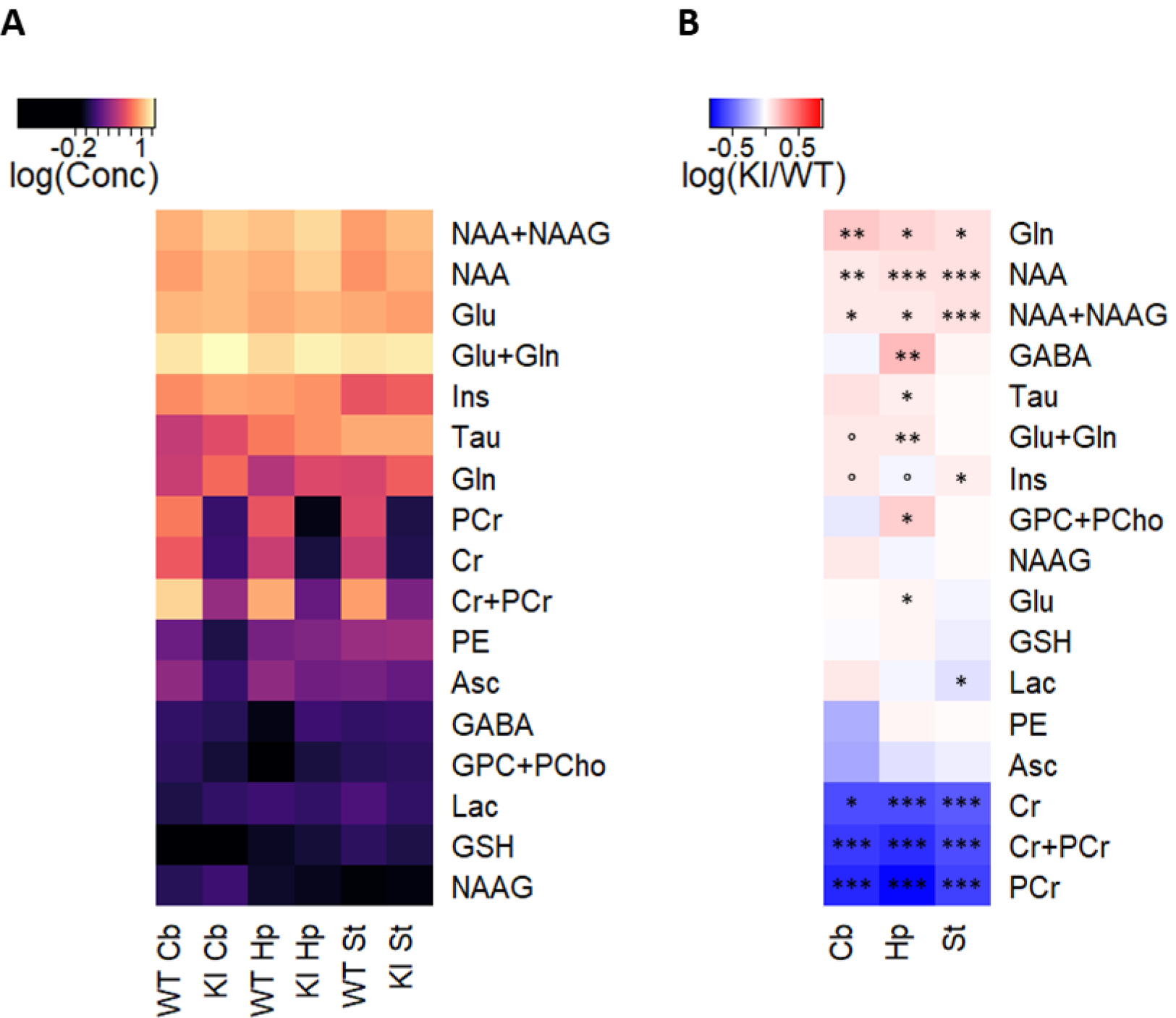
*Slc6a8*^*Y389C*>^ KI males present a systematic and significant increase in Gln, NAA and NAA+NAAG levels in all brain regions. (**A**) Logarithm (Log10) of concentration (in mmol/l) of ^1^H-MRS-measured metabolites in cerebellum (Cb), hippocampus (Hp) and striatum (St) from WT and KI males. (**B**) Logarithm of the KI concentrations normalized to WT (logarithm of the ratio between KI and WT concentrations). Blue colors indicate a reduction and red colors indicate an increase of concentrations with respect to the WT values. 3-4 WT and 4-5 KI animals; ° p<0.10; * p<0.05; ** p<0.01, *** p<0.001 (two-tail t-tests except for GABA in Cb, NAA+NAAG in Hp and PE in St, in which Mann-Whitney tests were used).

CTD patients present cognitive deficiency and behavioral troubles such as autistic-like behaviors or attention deficits. In order to analyze if our *Slc6a8*^*Y389C*^ rat line is a good model for neurological symptoms of CTD, we performed several behavioral tests in relation with those symptoms.

### *Slc6a8*^*xY389C/y*^ KI males show similar performance than WT ones on NOR and Y-maze for spatial memory tasks

To assess cognitive deficiency-like symptoms, we evaluated learning and memory performance in two different tasks. Y-maze test evaluates short-term spatial memory while Novel Object Recognition (NOR) test measures declarative memory. In Y-maze, both recognition index and total exploration time of the arms were similar between genotypes (**Figure 6 A and B**, respectively). In NOR, *Slc6a8*^*xY389C/y*^ KI males showed similar recognition index than WT littermates (**Figure 6 C**), although total exploration time of the objects was slightly higher in the testing phase (**Figure 6 D**) and significantly higher in the learning phase (+16% from WT, p=0.043). This increase in exploration time without any improvement in learning and memory performance may indicate attention deficits and/or impairment in working memory capacity.

**Figure 6:**
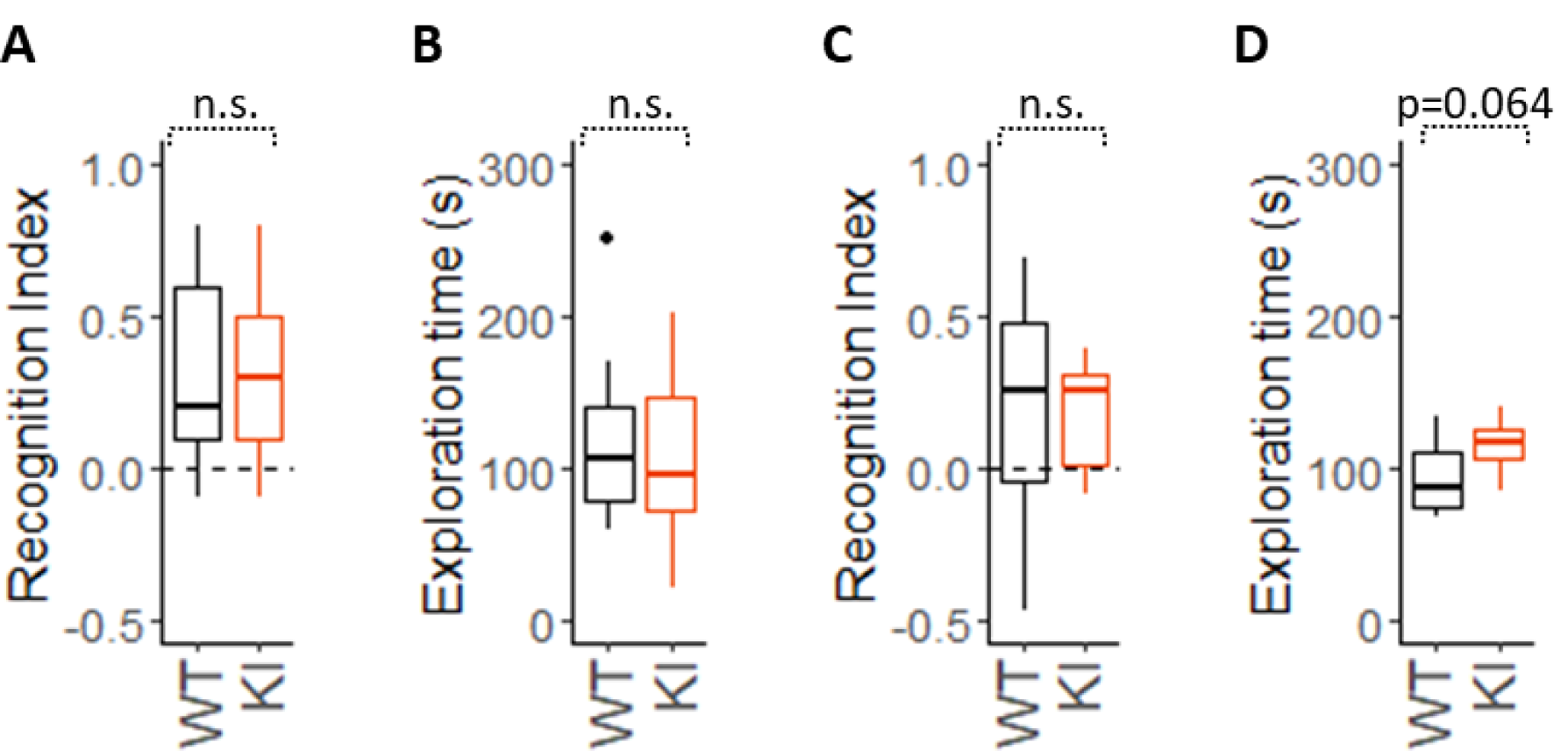
*Slc6a8*^*xY389C/y*^ KI males did not present differences in Y-maze (spatial memory) or NOR (declarative memory). **A-B**: Y-maze test showed no differences in recognition index (**A**) or total exploration time of both arms (**B**) between genotypes (p=0.94 and p=0.80, respectively). **C-D**: NOR test recognition index (**C**) was similar between genotypes (p=0.97), although total exploration time (in seconds) (**D**) was slightly higher in KI males (p=0.064). 9 WT + 9 KI males were used for Y-maze test, 9 WT + 9 KI for NOR test; p-values were estimated with two-tail t-tests. n.s.: not significant.

### *Slc6a8*^*xY389C/y*^ KI males present deficits in working memory

Working memory affects learning and memory ^32^. Additionally, to assess attention deficit in rodents, a task evaluating working memory is typically used. Thus, we carried out Y-maze for spontaneous alternation test in a different cohort of animals. Percentage of alternation index was used to compare working memory performance between genotypes and each genotype with the value of a random exploration (33.3% of alternation). While there were no differences in the number of entries between genotypes (**Figure 7 A**), the percentage of alternation for *Slc6a8*^*xY389C/y*^ KI males was significantly reduced in comparison with that of WT males, having similar value than the expected by chance (**Figure 7 B**). This result shows impaired working memory in KI males.

**Figure 7:**
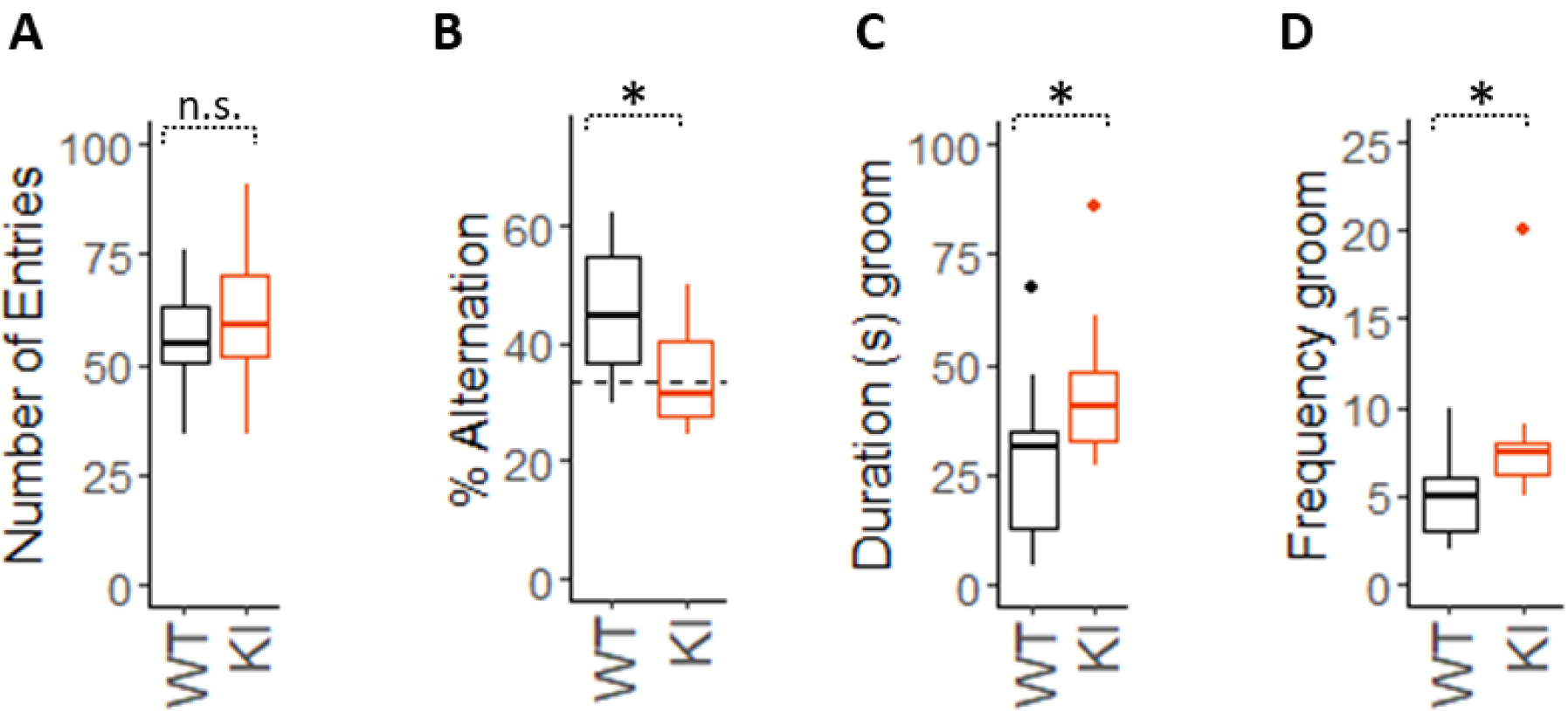
*Slc6a8*^*xY389C/y*^ KI males showed impaired working memory in Y-maze for spontaneous alternation, and more grooming behavior. **A-B:** Boxplots from Y-maze for spontaneous alternation test. (**A**) WT and KI males had no differences in the number of entries (p=0.57). (**B**) KI males showed a significant reduction in % of alternations when compared with WT littermates (p=0.039), being not higher than the value expected by chance (p=0.39 one-tail t-test) while % of alternations in WT males were significantly higher (p=0.010 one-tail t-test). (**C**) KI males showed more time doing grooming (p=0.044) and (**D**) with more frequency than WT littermates (p=0.028, Mann-Whitney test). 8 WT and 10 KI males were used for Y-maze for spontaneous alternations, 10 WT and 10 KI for grooming behavior. By default, p-values were estimated with two-tail t-test. n.s.: not significant; *: p<0.05.

### *Slc6a8*^*xY389C/y*^ KI males present more repetitive self-stimulating behavior

To analyze whether *Slc6a8*^*xY389C/y*^ KI males present autistic-like behavior, we decided to evaluate the presence of repetitive self-stimulating behaviors like grooming. In a period of 10 min in an open field, KI males spent significantly more time doing grooming than WT males, and with higher frequency (**Figure 7 C and D**, respectively).

### *Slc6a8*^*xY389C/y*^ KI males show a tendency in decreased sociability

Decreased sociability is one core feature of autism spectrum disorders. We used Social Preference test (SP) to evaluate the general sociability and Social Memory test (SM) to evaluate the interest in social novelty. For this analysis, we used the cumulative duration of the rat exploring an item (juvenile, object, new or familiar juvenile, **Figure 8 A, B**), the number of the visits for each item (frequency, **Figure 8 C**) and the decision of the animals on what item was explored at the first place (**Figure 8 D**). As expected, WT males presented a significant preference for the juvenile in the SP paradigm, by showing significantly more time exploring the juvenile instead of the object (p<0.001 two-tail t-test, **Figure 8 A** black boxplots). *Slc6a8*^*xY389C/y*^ KI males also showed social preference over the object (p<0.001 two-tail t-test, **Figure 8 A** orange boxplots), although spending a little less time with the juveniles in comparison with their WT littermates.

**Figure 8:**
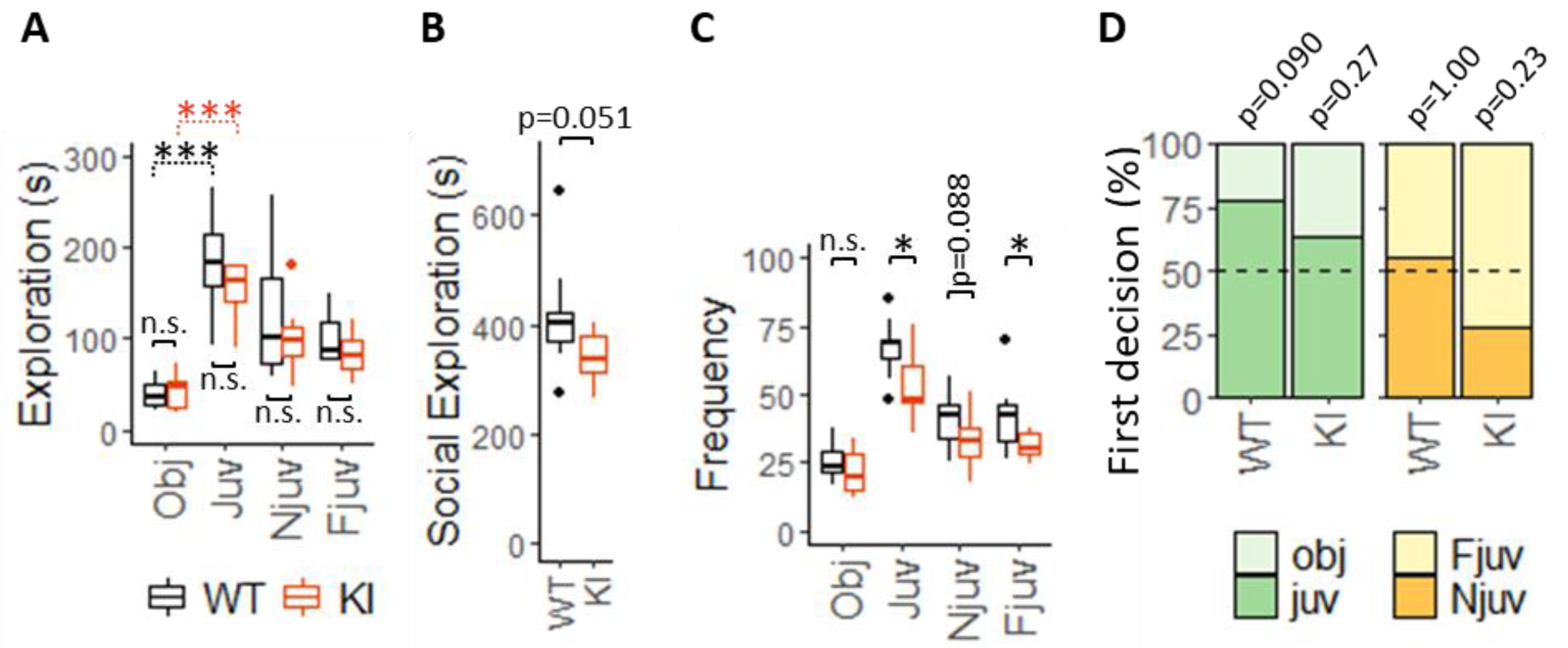
*Slc6a8*^*xY389C/y*^ KI males show a systematic tendency in decreased sociability. (**A**) Total exploration time (in seconds) for each item from WT and KI males (black and orange boxplots, respectively). Note that both WT and KI males explored significantly more the juvenile than the object in the social preference phase. Note also the tendency (although no significant) of KI males, in comparison with their WT littermates, to take more time exploring the object and less time all juveniles in all phases of the test. (**B**) Sum of the total exploration time (in seconds) of all the juveniles in all the SP-SM test is strongly reduced in KI males. (**C**) Number of visits (frequency) to each item for WT and KI males. Note the general significant decrease in the number of visits towards the juveniles in KI males. P-values in A-C were estimated with two-tail t-test. (**D**) Percentage of animals who went firstly to a certain item per each phase. In social preference, the number of WT males visiting the juvenile in the first place was marginally significant over the random expectation (dashed line, p=0.090 one-sided exact binomial test), while the number of KI males was reduced and near to the level expected by chance (p=0.27 one-sided exact binomial test). In social memory phase, while WT males’ preference to visit firstly the new or the familiar juvenile was random (p=1.00 two-sided exact binomial test), KI males tended to avoid the new juvenile in the first place (p=0.23 two-sided exact binomial test). 9 WT + 10 KI for both SP and SM phases. Obj, Juv, Njuv and Fjuv: object, juvenile, new and familiar juveniles respectively. *: p<0.05; ***: p<0.001.

Looking at the two phases together, *Slc6a8*^*xY389C/y*^ KI males systematically presented a tendency in reducing the cumulative duration exploring the juveniles (around −17.5% less than WT males) but not the object (+7.6%). These little changes of KI males resulted in a nearly significant reduction of the social exploration time along the two phases (**Figure 8 B**, p=0.051 two-tail t-test). Furthermore, the number of visits of KI rats towards the juveniles across the two phases was in general significantly lower than those of WT littermates but similar towards the object (**Figure 8 C**). As expected, we also observed that there were more WT animals going to explore firstly the juvenile (7 out of 9) instead of the object, being marginally significant with respect to the random expectation (p=0.090 one-sided exact binomial test, **Figure 8 D** green panel). In contrast, a lower percentage of KI animals decided to visit the juvenile in the first place (7 out of 11), being not significantly higher than the number expected by chance (p=0.27 one-sided exact binomial test, **Figure 8 D** green panel). In summary, *Slc6a8*^*xY389C/y*^ KI males presented social preference but with a systematic pattern of reduced sociability when compared to their WT littermates.

Regarding the social novelty paradigm, WT and *Slc6a8*^*xY389C/y*^ KI males spent similar time with the new and the familiar juveniles, showing no differences between the two genotypes (**Figure 8 A**). Interestingly, while the number of WT males exploring firstly the new juvenile was similar than the number expected by chance (5 out of 9 rats, p=1.00 two-sided exact binomial test, **Figure 8 C** yellow panel), the number of KI males tended to be lower (3 out of 11 rats, p=0.23 two-sided exact binomial test). These data may point towards an aversion of KI males in establishing the first contact with a social novelty.

## DISCUSION

We present here the first analysis of a new *in vivo* model of CTD, the *Slc6a8*^*Y389C*^ knock-in rat holding an identical patient’s missense mutation abolishing CrT activity ^17^. *Slc6a8*^*xY389C/y*^ KI rat males are brain creatine deficient, and exhibit increased Cr/Crn urine ratio as well as neurological symptoms similar to those of CTD patients, such as cognitive deficiency and some autistic features.

Using similar techniques as in the clinic, we first analyzed the brain metabolic profile of *Slc6a8*^*xY389C/y*^ KI rat males by ^1^H-MRS to check whether they present the same pathognomonic signature as CTD patients. All the *Slc6a8*^*xY389C/y*^ KI rat males analyzed showed the specific markers of CCDS and CTD: a strong reduction of Cr levels in the brain and a strong increase in urinary Cr/Crn ratio ^17^. These results validate the *Slc6a8*^*Y389C*^ rat line as a plausible model of CTD, being the first model of ubiquitous CrT deficiency confirming the two known biochemical signatures of CTD.

The increase in urine Cr levels reveals a functional deficiency in the reuptake of Cr from the primary urine, which is normally done at the renal proximal tubular cells by CrT ^33^. Given that plasma Cr is being filtered and lost in the urine due to CrT deficiency, we could expect that Cr levels in plasma were diminished and those of GAA, the precursor of Cr, were increased in an attempt to compensate Cr levels by AGAT feedback loop of up-regulation due to Cr deficiency (for this mechanism in periphery ^2,34,35^ or for CNS ^36^). Indeed, a tendency to decreased Cr levels and significant increased GAA were observed in the plasma of *Slc6a8*^*xY389C/y*^ KI males. This finding is in agreement with the decreased levels of Cr also observed in the ubiquitous KO mouse males with exons 2-4 deleted ^16^. These data contrast however with the low number of patients with available measurements of Cr and GAA in plasma showing normal or mildly elevated Cr levels as well as normal GAA ^7,17^. Further studies are needed to unravel the possible causes of these differences between human and rodents.

Furthermore, we present the first measurement of the GAA/Crn ratio in urine of an *in vivo* CTD model. This ratio was strongly and significantly elevated in *Slc6a8*^*xY389C/y*^ KI males, suggesting that GAA/Crn ratio in urine may also be a valid biochemical marker for CTD when is found together with increased Cr/Crn ratio. This result might help to bring attention towards a CTD diagnosis in patients with suspected GAMT-D (where increased GAA levels in urine are characteristic, although with a low or slightly low Cr/Crn ratio ^7^). New data from patients will help to confirm whether this double marker would be useful for CTD diagnosis. Whether GAA is also re-absorbed from primary urine in proximal tubular cells of kidney through transport by CrT (as we have shown in brain cells ^37^) remains to be determined. Indeed, while Cr was decreased in plasma and increased in urine of *Slc6a8*^*xY389C/y*^ KI males demonstrating CrT deficiency, GAA was increased in plasma but normal with a tendency to increase in urine.

Consistent with CTD patients ^17^ and all the ubiquitous CrT KO mouse models ^12,14,16^, *Slc6a8*^*xY389C/y*^ KI males presented lower body weight. We observed this difference already after weaning, but taking into account the degree of significance, it is likely that it extends from previous time points. This body weight gain issue is in agreement with reported early signs of CTD ^38^.

Although Cr levels in plasma and in brain were lower in *Slc6a8*^*xY389C/y*^ KI males than in their WT littermates, Cr levels in the CSF were mildly higher or similar to those of WT ones. This is consistent with what was observed in CTD patients ^7^, confirming that CSF does not always reflect what is happening in brain parenchyma, and that CSF can also be a useful diagnostic sample for the differential diagnosis between CTD and GAMT deficiency, in which Cr is decreased ^7^. Interestingly, while WT males showed higher levels of Cr in plasma than in CSF, *Slc6a8*^*xY389C/y*^ KI males exhibited similar levels in both liquids (60-70 μmol/l). Altogether, these data support the fact that the Cr pool in CNS does not fully depend on the peripheral one, owing to the low permeability of blood-brain barrier for Cr and to brain endogenous synthesis of Cr ^9,37,39,40^.

In every brain region analyzed, the strongly decreased Cr levels in brain parenchyma from *Slc6a8*^*xY389C/y*^ KI males were accompanied by significant increases in Gln, NAA and NAA+NAAG. Given that these molecules are important osmolytes in CNS ^41-43^, and that the magnitudes of both changes tend to be similar, it might hint at an osmolyte compensation mechanism by Gln, NAA and/or NAAG for the brain Cr deficiency. Other metabolites with a role in osmoregulation are taurine, myo-inositol (Ins) or choline ^44-46^. Interestingly, the concentration of these metabolites change in some of the regions analyzed as well (see below). We have recently shown a similar potential mechanism of osmolytes compensation involving the same molecules in another pathological condition of CNS (hepatic encephalopathy ^24^). As Gln, NAA and NAAG play other important roles in CNS, a line for future research is to establish whether their changes only compensate Cr deficiency osmotically, or are truly pathological.

Other metabolites changed differentially depending on the brain region analyzed. In hippocampus, neurotransmitters such as GABA, Glu and Tau increased together with total choline, a precursor for another neurotransmitter (acetylcholine), membrane phospholipids and, (via betaine) the methyl group donor *S*-adenosylmethionine which is involved in the synthesis of Cr from GAA ^47^. Gln is a precursor for Glu, which is a precursor for GABA. Transporters for GABA and Tau can transport Cr or GAA ^39^. It would thus be possible that alterations in Cr and/or GAA levels might affect GABA and Tau homeostasis and their concentrations. As described above Tau and total choline in hippocampus might also participate to an osmolyte compensation mechanism under Cr deficiency. Overall, these metabolic changes may suggest that endogenous Cr synthesis may also be affected, together with disturbances in neurotransmission in the hippocampus of *Slc6a8*^*xY389C/y*^ KI males. Hippocampus is involved in motivation, emotion, learning and memory, and these changes could affect its functions and also promote epilepsy in CTD patients. In striatum, *Slc6a8*^*xY389C/y*^ KI males showed an increase in Ins, osmolyte also involved in many other cellular processes. The decrease in lactate might reflect a change in striatal energy metabolism.

These alterations in brain metabolites correlate with changes in behavior and cognition. In comparison with their WT littermates, *Slc6a8*^*xY389C/y*^ KI males showed similar learning and memory performance in simple tasks about recognition and spatial memory (NOR and Y-maze for spatial memory tests) but failed in a working memory task (Y-maze for spontaneous alternation). Higher exploration times of objects in both phases of NOR test without any advantage in declarative memory might reflect a deficit in cognition related with attention and working memory as well. These negative results in learning and memory performance contrast with the data from the ubiquitous CrT KO mice with exons 2-4 and 5-7 deleted, where deficits were present although with milder phenotype in NOR test around the same age in the last one ^13,16^. Interestingly, these negative results are consistent with data from CTD patients, as they are able to remember people, objects or locations but fail in the performance of more complex daily tasks and learning new abilities ^17,38^.

On the other hand, working memory deficits of KI males are in agreement with the ubiquitous CrT KO with exons 5-7 deleted ^13^ and with the presence of attention deficits observed in patients ^17^. However, not all CTD patients, including the one sharing the c.1166A<G; p.(Tyr389Cys) mutation with KI males, are diagnosed with attention deficits ^17^ (Van de Kamp personal communication).

A lot of evidence about working memory supports the relationship with learning and memory processes ^32^ as well as the significant impact on the performance of daily tasks ^48^. Working memory capacity is a good predictor of learning outcomes ^49^ and general intelligence ^50^. Overall, our data may suggest that impairment in working memory, rather than deficits in declarative memory *per se*, is one of the primary components in the intellectual disability from CTD patients. Such hypothesis could open new avenues for alternative therapies to support working memory in order to improve learning in patients. Further studies are needed to address these questions.

Besides intellectual disability and attention deficits, autistic-like behavior is another common symptom in CTD patients. We evaluated the presence of two core features of autism spectrum disorder: repetitive self-stimulating behaviors (like grooming for rats) and sociability. *Slc6a8*^*xY389C/y*^ KI males spent significantly more time in grooming and with more frequency, consistent with another CTD model ^13^. Additionally, KI males appeared to be social but less than their WT littermates. This less social tendency is seen in the systematic pattern of spending less time with less frequency exploring the juveniles in all phases in contrast to exploring the object, and by having less tendency to first visit the juvenile instead of the object. Additionally, while WT males showed no preference to first visit any of the juveniles, KI males showed slightly more preference towards the familiar juvenile in the first place. This result might reflect an aversion to establish the first contact with a social novelty, which is another feature of sociability present in autism. Interestingly, these results are in line with the autistic phenotype of the CTD patient sharing the same missense mutation as *Slc6a8*^*xY389C/y*^ KI males ^17^ (Van de Kamp, personal communication). Importantly, the *Slc6a8*^*Y389C*^ rat line is the first animal model of CTD showing a difference in sociability. Further studies will be needed to unravel the nature of this difference.

In summary, this new *Slc6a8*^*Y389C*^ rat line is the first CTD model generated so far with one of the single nucleotide mutations described in CTD patients. *Slc6a8*^*xY389C/y*^ male rats present characteristic features of CTD and reveal the *Slc6a8*^*Y389C*^ rat line as a valuable model of CTD, fairly enriching the spectrum of *in vivo* CTD models. Additionally, we have shown new data confirming what has been shown in patients and highlighting common aspects of CTD pathology. Future work will aim at further characterizing the consequences of CTD in the *Slc6a8*^*Y389C*^ rat line both in periphery and in CNS (in particular to better understand the mechanisms underlying metabolic changes and disturbances of brain development and behavior), and to use this *in vivo* model to develop new strategies of treatment for CTD.

## Acknowledgements

This work was supported by the Swiss National Science Foundation (grant n° 31003A-175778). We thank Marc Loup, Dario Sessa, Martine Vaglio and Ana Versace for excellent technical help, Isabelle Guillot de Suduiraut for helpful discussion with behavioral tests, and Alberto Pascual-Garcia for statistical and computational advice. The authors declare no competing financial interests.

## Data availability statement

All data generated or analyzed during this study are included in this published article.

## Bibliography

1 Hanna-El-Daher, L. & Braissant, O. Creatine synthesis and exchanges between brain cells: What can be learned from human creatine deficiencies and various experimental models? Amino acids 48, 1877–1895, doi: 10.1007/s00726-016-2189-0 (2016).

2 Wyss, M. & Kaddurah-Daouk, R. Creatine and creatinine metabolism. Physiological reviews 80, 1107–1213, doi: 10.1152/physrev.2000.80.3.1107 (2000).

3 Item, C. B. et al. Arginine:glycine amidinotransferase deficiency: the third inborn error of creatine metabolism in humans. American journal of human genetics 69, 1127–1133, doi: 10.1086/323765 (2001).

4 Schulze, A. & Braissant, O. in Pediatric endocrinology and inborn errors of metabolism (eds Kyriakie Sarafoglou, Georg F Hoffmann, & Karl S Roth) 181–190 (McGraw Hill, 2017).

5 Salomons, G. S. et al. X-linked creatine-transporter gene (SLC6A8) defect: a new creatine-deficiency syndrome. The American Journal of Human Genetics 68, 1497–1500 (2001).

6 Stöckler, S. et al. Creatine deficiency in the brain: a new, treatable inborn error of metabolism. Pediatric Research 36, 409–413 (1994).

7 Van de Kamp, J. M., Mancini, G. M. & Salomons, G. S. X-linked creatine transporter deficiency: clinical aspects and pathophysiology. Journal of inherited metabolic disease 37, 715–733, doi: 10.1007/s10545-014-9713-8 (2014).

8 Rackayova, V., Cudalbu, C., Pouwels, P. J. W. & Braissant, O. Creatine in the central nervous system: From magnetic resonance spectroscopy to creatine deficiencies. Analytical biochemistry 529, 144–157, doi: 10.1016/j.ab.2016.11.007 (2017).

9 Abdulla, Z. I., Pennington, J. L., Gutierrez, A. & Skelton, M. R. Creatine transporter knockout mice (Slc6a8) show increases in serotonin-related proteins and are resilient to learned helplessness. Behavioural brain research 377, 112254, doi: 10.1016/j.bbr.2019.112254 (2020).

10 Udobi, K. C. et al. Deletion of the creatine transporter gene in neonatal, but not adult, mice leads to cognitive deficits. Journal of inherited metabolic disease 42, 966–974, doi: 10.1002/jimd.12137 (2019).

11 Udobi, K. C. et al. Cognitive deficits and increases in creatine precursors in a brain-specific knockout of the creatine transporter gene Slc6a8. Genes, brain, and behavior 17, e12461, doi: 10.1111/gbb.12461 (2018).

12 Stockebrand, M. et al. A Mouse Model of Creatine Transporter Deficiency Reveals Impaired Motor Function and Muscle Energy Metabolism. Frontiers in physiology 9, 773, doi: 10.3389/fphys.2018.00773 (2018).

13 Baroncelli, L. et al. A mouse model for creatine transporter deficiency reveals early onset cognitive impairment and neuropathology associated with brain aging. Hum Mol Genet 25, 4186–4200, doi: 10.1093/hmg/ddw252 (2016).

14 Baroncelli, L. et al. A novel mouse model of creatine transporter deficiency. F1000Research 3, 228, doi: 10.12688/f1000research.5369.1 (2014).

15 Kurosawa, Y. et al. Cyclocreatine treatment improves cognition in mice with creatine transporter deficiency. The Journal of clinical investigation 122, 2837–2846, doi: 10.1172/jci59373 (2012).

16 Skelton, M. R. et al. Creatine transporter (CrT; Slc6a8) knockout mice as a model of human CrT deficiency. PLoS One 6, e16187, doi: 10.1371/journal.pone.0016187 (2011).

17 Van de Kamp, J. M. et al. Phenotype and genotype in 101 males with X-linked creatine transporter deficiency. J Med Genet 50, 463–472, doi: 10.1136/jmedgenet-2013-101658 (2013).

18 Molinaro, A. et al. A Nervous System-Specific Model of Creatine Transporter Deficiency Recapitulates the Cognitive Endophenotype of the Disease: a Longitudinal Study. Scientific reports 9, 62, doi: 10.1038/s41598-018-37303-1 (2019).

19 Abdulla, Z. I. et al. Deletion of the Creatine Transporter (Slc6a8) in Dopaminergic Neurons Leads to Hyperactivity in Mice. Journal of molecular neuroscience : MN 70, 102–111, doi: 10.1007/s12031-019-01405-w (2020).

20 Drummond, E. & Wisniewski, T. Alzheimer’s disease: experimental models and reality. Acta neuropathologica 133, 155–175, doi: 10.1007/s00401-016-1662-x (2017).

21 Do Carmo, S. & Cuello, A. C. Modeling Alzheimer’s disease in transgenic rats. Molecular neurodegeneration 8, 37, doi: 10.1186/1750-1326-8-37 (2013).

22 Ellenbroek, B. & Youn, J. Rodent models in neuroscience research: is it a rat race? Disease models & mechanisms 9, 1079–1087, doi: 10.1242/dmm.026120 (2016).

23 Gruetter, R. Automatic, localized in vivo adjustment of all first-and second-order shim coils. Magnetic resonance in medicine 29, 804–811 (1993).

24 Braissant, O. et al. Longitudinal neurometabolic changes in the hippocampus of a rat model of chronic hepatic encephalopathy. Journal of hepatology 71, 505–515 (2019).

25 Mlynárik, V., Gambarota, G., Frenkel, H. & Gruetter, R. Localized short-echo-time proton MR spectroscopy with full signal-intensity acquisition. Magnetic Resonance in Medicine: An Official Journal of the International Society for Magnetic Resonance in Medicine 56, 965–970 (2006).

26 Provencher, S. W. Automatic quantitation of localized in vivo 1H spectra with LCModel. NMR in Biomedicine: An International Journal Devoted to the Development and Application of Magnetic Resonance In Vivo 14, 260–264 (2001).

27 Cudalbu, C., Mlynárik, V. & Gruetter, R. Handling macromolecule signals in the quantification of the neurochemical profile. Journal of Alzheimer’s Disease 31, S101–S115 (2012).

28 Team, R. C. R: A Language and Environment for Statistical Computing (Version 3.5. 1, R Foundation for Statistical Computing, Vienna, Austria, 2018). https://www.R-project.org/. (2018).

29 Wickham, H. ggplot2: elegant graphics for data analysis. (springer, 2016).

30 Warnes, M. G. R., Bolker, B., Bonebakker, L., Gentleman, R. & Huber, W. Package ‘gplots’. Various R Programming Tools for Plotting Data (2016).

31 Calhoun, P. Exact: Unconditional Exact Test. R package version 2.0.. https://CRAN.R-project.org/package=Exact (2019).

32 Cowan, N. & Alloway, T. The development of working memory. The development of memory in childhood, 163–199 (1997).

33 García-Delgado, M., Peral, M. J., Cano, M., Calonge, M. L. & Ilundáin, A. A. Creatine transport in brush-border membrane vesicles isolated from rat kidney cortex. Journal of the American Society of Nephrology : JASN 12, 1819–1825 (2001).

34 Guthmiller, P., Van Pilsum, J., Boen, J. R. & McGuire, D. M. Cloning and sequencing of rat kidney L-arginine: glycine amidinotransferase. Studies on the mechanism of regulation by growth hormone and creatine. Journal of Biological Chemistry 269, 17556–17560 (1994).

35 McGuire, D. M., Gross, M. D., Van Pilsum, J. & Towle, H. C. Repression of rat kidney L-arginine: glycine amidinotransferase synthesis by creatine at a pretranslational level. Journal of Biological Chemistry 259, 12034–12038 (1984).

36 Braissant, O. et al. Ammonium alters creatine transport and synthesis in a 3D culture of developing brain cells, resulting in secondary cerebral creatine deficiency. European Journal of Neuroscience 27, 1673–1685 (2008).

37 Braissant, O., Béard, E., Torrent, C. & Henry, H. Dissociation of AGAT, GAMT and SLC6A8 in CNS: relevance to creatine deficiency syndromes. Neurobiology of disease 37, 423–433 (2010).

38 Miller, J. S. et al. Early Indicators of Creatine Transporter Deficiency. The Journal of pediatrics 206, 283–285, doi: 10.1016/j.jpeds.2018.11.008 (2019).

39 Braissant, O. Creatine and guanidinoacetate transport at blood-brain and blood-cerebrospinal fluid barriers. Journal of inherited metabolic disease 35, 655–664 (2012).

40 Braissant, O., Henry, H., Loup, M., Eilers, B. & Bachmann, C. Endogenous synthesis and transport of creatine in the rat brain: an in situ hybridization study. Molecular brain research 86, 193–201 (2001).

41 Albrecht, J., Sonnewald, U., Waagepetersen, H. S. & Schousboe, A. Glutamine in the central nervous system: function and dysfunction. Frontiers in bioscience : a journal and virtual library 12, 332–343, doi: 10.2741/2067 (2007).

42 Moffett, J. R., Ross, B., Arun, P., Madhavarao, C. N. & Namboodiri, A. M. N-Acetylaspartate in the CNS: from neurodiagnostics to neurobiology. Prog Neurobiol 81, 89–131, doi: 10.1016/j.pneurobio.2006.12.003 (2007).

43 Neale, J. H., Bzdega, T. & Wroblewska, B. N-Acetylaspartylglutamate: the most abundant peptide neurotransmitter in the mammalian central nervous system. J Neurochem 75, 443–452, doi: 10.1046/j.1471-4159.2000.0750443.x (2000).

44 Waldegger, S., Matskevitch, J., Busch, G. L. & Lang, F. Introduction to cell volume regulatory mechanisms. Contributions to nephrology 123, 1–7, doi: 10.1159/000059919 (1998).

45 Bauernschmitt, H. & Kinne, R. Metabolism of the ‘organic osmolyte’glycerophosphorylcholine in isolated rat inner medullary collecting duct cells. Pathways for synthesis and degradation. Biochimica et Biophysica Acta (BBA)-Biomembranes 1148, 331–341 (1993).

46 Oja, S. S. & Saransaari, P. Significance of Taurine in the Brain. Advances in experimental medicine and biology 975 Pt 1, 89–94, doi: 10.1007/978-94-024-1079-2_8 (2017).

47 Bekdash, R. A. Choline, the brain and neurodegeneration: insights from epigenetics. Frontiers in bioscience (Landmark edition) 23, 1113–1143, doi: 10.2741/4636 (2018).

48 Humphreys, G. W., Forde, E. M. & Francis, D. The organization of sequential actions. Control of cognitive processes: Attention and performance XVIII, 425–472 (2000).

49 Alloway, T. P. & Alloway, R. Working memory: Is it the new IQ? Nature Precedings, doi: 10.1038/npre.2008.2343.1 (2008).

50 Conway, A. R., Cowan, N., Bunting, M. F., Therriault, D. J. & Minkoff, S. R. A latent variable analysis of working memory capacity, short-term memory capacity, processing speed, and general fluid intelligence. Intelligence 30, 163–183 (2002).

